# A humanized ossicle model of myelofibrosis reveals THPO-driven fibrosis, osteosclerosis and SPP1-dependent microenvironmental remodeling

**DOI:** 10.64898/2026.03.12.711163

**Authors:** Hongzhe Li, Alba Lillo Sierras, Rong Fan, Michaela Oeller, Katharina Schallmoser, Anne Hultquist, Stefan Scheding

## Abstract

Myelofibrosis (MF) is the most severe myeloproliferative neoplasm. Current therapies – except for allogeneic stem cell transplantation – are largely supportive which highlights the need for improved disease models and novel therapeutic targets.

Here, we established a humanized MF model by transplanting thrombopoietin (THPO)-overexpressing human bone marrow CD34□ cells into humanized bone marrow ossicles generated in immunodeficient NSG mice. THPO overexpression induced progressive reticulin fibrosis in vivo, accompanied by myeloid skewing, increased megakaryocyte clustering, and redistribution of human hematopoietic cells to murine spleen and femur, consistent with extramedullary hematopoiesis. THPO-driven ossicles also exhibited features of osteosclerosis, including increased trabecular bone and osteoid formation, indicating active pathological remodeling of the niche.

Mechanistically, fibrosis was associated with increased SPP1/OPN expression, which was also observed in bone marrow biopsies from MF patients. Importantly, in vivo neutralization of SPP1 attenuated myeloid skewing, reduced megakaryocyte expansion, and decreased fibrosis severity, highlighting SPP1-driven niche remodeling as a potential therapeutic target in MF. This humanized MF model thus provides a translationally relevant platform to dissect microenvironment-driven MF pathogenesis and evaluate targeted therapies.

## Introduction

Myelofibrosis (MF) is a chronic myeloproliferative neoplasm (MPN) characterized by clonal expansion of hematopoietic stem and progenitor cells (HSPCs), bone marrow (BM) fibrosis, extramedullary hematopoiesis, progressive cytopenias and osteosclerosis [1, 2]. MF is the most aggressive form among the BCR-ABL1-negative MPNs with a considerable risk for transformation to acute leukemia. MF can occur de novo as primary myelofibrosis (PMF) or evolve from essential thrombocythemia (ET) or polycythemia vera (PV) as post-ET and post-PV myelofibrosis, respectively. Driver mutations in Janus kinase 2 (JAK2), calreticulin (CALR), or myeloproliferative leukemia virus oncogene (MPL) are central to MF pathogenesis, leading to constitutive JAK-STAT signaling and dysregulated cytokine production, which drive abnormal megakaryopoiesis, stromal remodeling, and reticulin fiber formation [3].

Despite the introduction of JAK inhibitors such as ruxolitinib, current therapies for MF are largely supportive [4]. JAK inhibitors provide mainly symptomatic relief and reduce splenomegaly but have limited impact on bone marrow fibrosis and disease progression. Although a number of novel MF therapies are under development, allogeneic stem cell transplantation (allo-HSCT) is currently the only curative treatment. Allo-HSCT, however, is a viable option for only a minority of patients due to the higher age of the MF population and corresponding high transplant-related morbidity and mortality. Thus, there is an urgent need for the development of improved therapeutic strategies based on targeting key pathophysiological mechanism in MF.

Here, murine MF models, which are based on thrombopoietin (THPO) overexpression and MPN driver mutation expression, respectively, provide valuable insights into MF pathophysiology and have been widely used for MF drug development [5–10]. However, species-specific differences in cytokine signaling, megakaryocyte biology, and stromal responses limit their translational relevance.

We therefore aimed to establish a humanized xenotransplantation model that more closely reflects human MF pathology. We generated humanized BM tissues (ossicles) in immunodeficient NSG mice by subcutaneous transplantation of human BM stromal cells followed by intra-ossicle (i.o.) transplantation of THPO overexpressing human CD34^+^ BM cells to induce fibrosis (MF-ossicles). Our results demonstrate that the model reproduced key MF features and that novel anti-fibrosis targets could be identified and functionally validated in-vivo.

Thus, this novel MF-ossicle model provides a unique and physiologically relevant platform to study MF pathogenesis and to evaluate therapeutic responses, thereby offering valuable insights for the development of more effective treatments for MF.

## Materials and Methods

### Human bone marrow mononuclear cell isolation and stromal cell culture

Human bone marrow (BM) samples were obtained from consenting healthy adult donors at the Hematology Department, Skåne University Hospital, Lund, Sweden, by aspiration from the iliac crest, as described previously [11]. The procedure was approved by the Regional Ethics Review Board in Lund, Sweden.

In brief, 60□mL of BM were aspirated from each donor into heparin-containing syringes. Bone marrow mononuclear cells (BM-MNCs) were isolated by density gradient centrifugation using LSM 1077 Lymphocyte separation medium (PAA, Pasching, Austria). BM stromal cells (BM-MSCs) were culture-derived from BM-MNCs as described previously [12]. Briefly, BM-MNCs were resuspended in pre-warmed α-modified minimum essential medium (α-MEM; Sigma-Aldrich, St. Louis, MO, USA), supplemented with 10% pooled human platelet lysate (pHPL, Department of Transfusion Medicine, Paracelsus Medical University, Salzburg, Austria), and seeded into tissue culture vessels. After initial incubation, non-adherent cells were removed by washing with phosphate-buffered saline (PBS), and adherent BM stromal cells (MSCs) were subsequently expanded until the required cell number was reached.

### Human bone marrow biopsies

Formalin-fixed paraffin-embedded (FFPE) BM biopsies from PMF patients with fibrosis grades 2 (n=3) or controls (n=3) were analyzed. Control biopsies were diagnosed as normal by routine pathology. Patient characteristics are provided in Table S1. BM specimens were obtained from the Division of Pathology, Laboratory Medicine Skåne, Lund, Sweden. Five µm thick sections were cut from paraffin blocks and transferred to slides. The sampling of patient material and sample preparation was approved by the institutional review committee in Lund (Regionala Etikprövningsnämnden in Lund, approvals no. 2014/776 and 2018/86) and all procedures were performed following the Helsinki Declaration of 1975, as revised in 2008.

### Humanized ossicle formation and human hematopoietic cell transplantation

Passage 2-3 cultured human BM-MSCs were harvested by trypsinization (Gibco, Thermo Fisher Scientific, Waltham, MA, USA). A total of 2 × 10□ stromal cells were resuspended in 60 µL of pooled human platelet lysate (pHPL) and mixed with 240 µL of Matrigel-equivalent matrix (In Vitro Angiogenesis Assay Kit, Merck, Darmstadt, Germany) plus Endothelial Cell Growth supplement (ECGS, Discovery labware Inc, Bedford, MA, USA) to obtain a final volume of 300 µL per injection. The cell–matrix mixture was then injected subcutaneously into both flanks of 7–8-week-old immunodeficient NOD.Cg-Prkdc^scid^Il2rγ^tm1Wjl^/SzJ (NSG) mice (Jackson Laboratory, Bar Harbor, ME, USA) to generate humanized ossicles (two ossicles per mouse).

Beginning on day 3 after implantation, mice received daily subcutaneous injections of human parathyroid hormone [PTH (1–34), Sigma-Aldrich, St. Louis, MO, USA; 40 µg/kg body weight] into the dorsal neck fold for 28 days to promote bone formation. Eight weeks after stromal cell implantation, ossicle-bearing mice were subjected to sublethal irradiation (200 cGy). At least four hours after irradiation, 1–3 × 10□ human BM CD34□ cells—either non-transduced, lentiviral GFP-transduced, or lentiviral THPO-GFP-transduced—were injected directly into each ossicle (intraossicle, i.o.).

Generally, mice were euthanized eight weeks after CD34□ cell transplantation, and tissues including ossicles, spleen, femurs, and liver were collected for downstream analyses. For time course fibrosis analysis, selected groups of mice were analyzed at different time points after 4 - 10 weeks. All animal experiments were performed in accordance with institutional and national ethical guidelines and were approved by the Malmo/Lund ethics committee for animal research.

### Lentiviral transduction of BM CD34□ Cells

MACS-enriched human bone marrow CD34□ cells were cultured overnight in StemSpan SFEM (StemCell Technologies, Vancouver, BC, Canada) supplemented with recombinant human SCF, THPO, and FLT3L (each at 100 ng/mL; PeproTech, Rocky Hill, NJ, USA). The following day, CD34□ cells were transduced with lentiviral particles encoding either THPO-GFP or GFP control at a multiplicity of infection (MOI) of 5. The lentiviral expression plasmids (THPO-GFP and GFP control) and the corresponding viral preparations were generated by the Cell and Gene Technologies Core Facility at the Lund Stem Cell Center, Lund, Sweden. Twenty-four hours after transduction, cells were collected, washed, resuspended in PBS, and used for in vivo transplantation.

### In vivo antibody administration

Anti-human SPP1/OPN antibody and corresponding isotype control antibody, respectively (both at 15 mg/kg body weight; Bioxcell, Lebanon, NH, USA) were administered via intraperitoneal injection. Treatment was initiated at week 4 and continued once weekly for two weeks. From week 6 onward, anti-human SPP1/OPN antibody was administered twice weekly until the experimental endpoint. Antibody doses and administration schedules were selected based on previously published studies [13, 14].

### Cell isolation from ossicles, femur, spleen, and liver

At indicated time points after intraossicle transplantation of human BM CD34□ cells, mice were euthanized and ossicles, spleens, femurs, and livers were collected for cell isolation.

#### Ossicles

Each ossicle was mechanically dissociated using a mortar and pestle, followed by enzymatic digestion in ossicle digestion buffer (Phosphate Buffered Saline, PBS, containing 5% Fetal Bovine Serum, FBS, 2 mg/mL collagenase II, and 4 mg/mL dispase II; 5 mL per ossicle) at 37 °C for 1 h with gentle agitation. The digested suspension was passed through a 100-µm cell strainer, and enzymatic activity was quenched by adding 35 mL of ice-cold PBS supplemented with 10% FBS. Cells were pelleted by centrifugation, washed twice with PBS + 10% FBS, and centrifuged again.

#### Spleen and liver

Tissues were mechanically dissociated by cutting them into small fragments, followed by gentle crushing and repeated pipetting in PBS to obtain single-cell suspensions.

#### Femur

After removal of both femoral heads, bone marrow was flushed from the diaphysis using a syringe and needle with 10 mL PBS + 5% FBS. The remaining femoral shafts and heads were then cut into small fragments and further dissociated using a mortar and pestle.

Cells isolated from ossicles, spleen, liver, and femur were subjected to red blood cell (RBC) lysis prior to antibody staining. RBC lysis was performed using 1× Pharm Lyse (BD Biosciences, Franklin Lakes, NJ, USA) according to the manufacturer’s instructions.

### Flow cytometry

Cells isolated from the different tissues were incubated in blocking buffer [Dulbecco’s PBS w/o Ca2+, Mg2+, 3.3 mg/ml human normal immunoglobubin (Gammanorm, Octapharm, Stockholm, Sweden), 1% FBS (Invitrogen, Carlsbad, CA, USA)], followed by staining with antibodies against human CD45, CD15, CD33, CD66b, CD19, CD41a, CD42b, CD34, CD56 and CD271. Antibodies are listed below. Gates were set according to the corresponding fluorescence-minus-one (FMO) controls and cells were analyzed on FACSymphony A1 cell analyser (BD Bioscience, Franklin Lakes, NJ, USA). Dead cells were excluded by DAPI (Merck) staining and doublets were excluded by gating on FSC-H versus FSC-W and SSC-H versus SSC-W.

### Antibodies

For FACS analysis, the following antibodies were used: hCD45-APC (clone HI30), hCD15-PE (clone HI98), hCD66b-PE (clone G10F5), hCD45-APC-Cy7 (clone 2D1) were from BD Pharmingen (San Jose, CA, USA), hCD19-PE-Cy5 (clone HIB19). hCD33-PE (clone P67.6), hCD41a-PE (clone HIP8), CD42b-APC (clone HIP1), hCD34-PerCP-Cy5.5 (clone 8G12) were from BD Biosciences (Franklin Lakes, NJ, USA). hCD271-APC (ME20.4-1.H4) was from Miltenyi Biotec (Bergisch Gladbach, Germany).

### Other methods

The details of the other methods used in this study, i.e. Hematoxylin and Eosin (H&E) staining, Gordon and Sweet’s reticulin staining, trichrome RGB staining, immunohistochemistry (IHC), immunofluorescence staining, and bone and megakaryocyte quantification are provided in the Supplementary information.

## Results

### Generation of a humanized bone marrow ossicles in NSG mice

To establish an ectopic humanized bone marrow (BM) niche capable of supporting human hematopoiesis, we adopted the approach described by Reinisch et al. [17] (Fig. 1A). Briefly, bone marrow-derived stromal cells from healthy donors were culture-expanded and injected subcutaneously into immunodeficient NSG mice together with an angiogenic hydrogel. Over time, this led to the formation of a bony ossicle (huOssicle) (Fig. 1B) containing functional bone marrow tissue with key anatomical features of human bone marrow including cortical bone, trabecular bone, adipocytes, fibroblastic stroma, and blood vessels (Fig. 1C). We also identified distinct periosteal and endosteal regions and that ossicle BM was infiltrated by murine hematopoietic cells (Fig. 1C). Importantly, staining for human-specific mitochondria confirmed the human origin of osteocytes and stromal elements (Fig. 1D).

**Figure 1.**
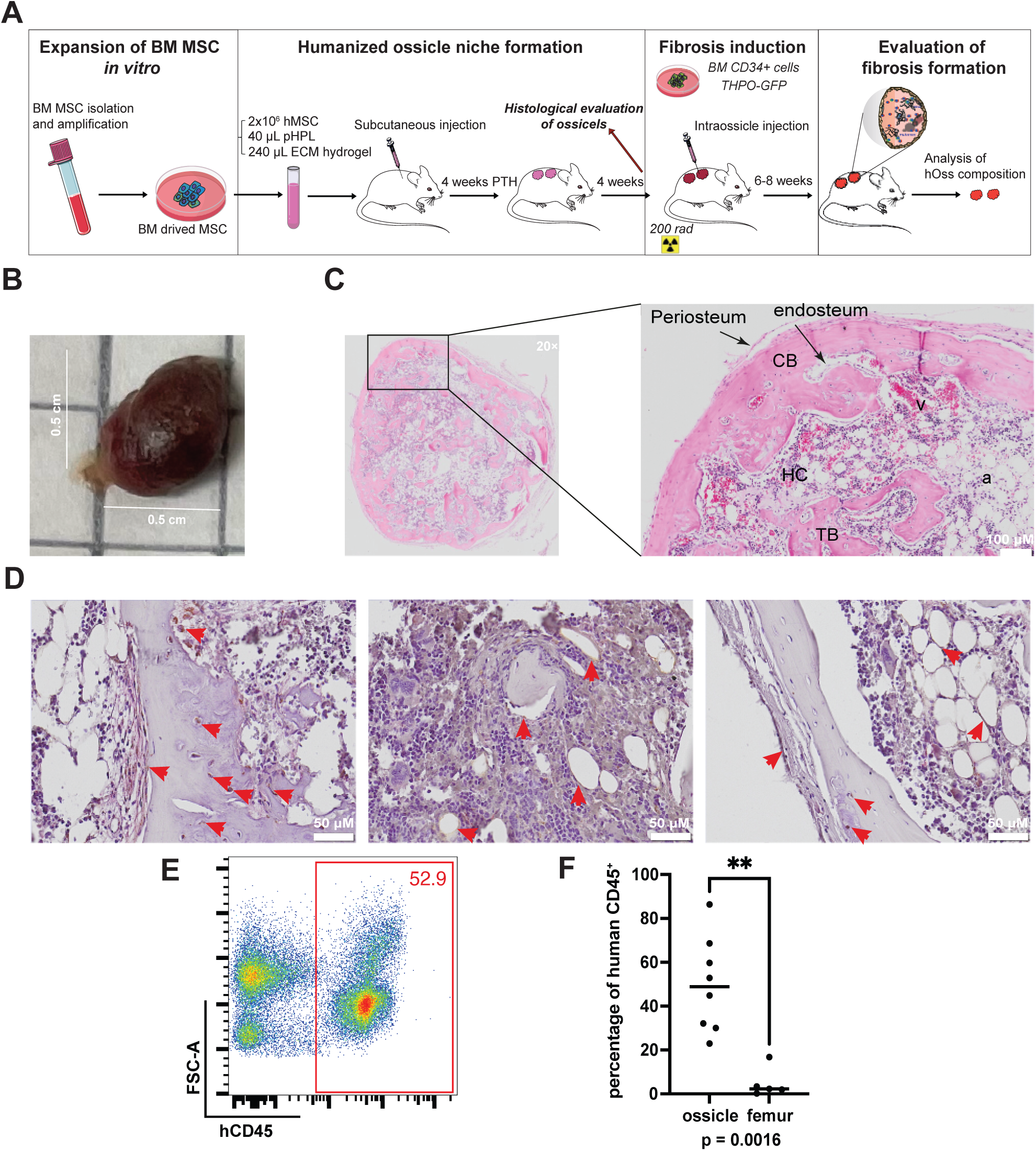
Generation of a humanized bone marrow ossicles in NSG mice. (A) Schematic representation of the experimental design for generating humanized ossicles and inducing myelofibrosis. BM, bone marrow; MSC, marrow stromal cells; pHPL, pooled human platelet lysate; PTH, parathyroid hormone; THPO, thrombopoietin; huMF, human myelofibrosis; hOss, humanized ossicle; ECM, extracellular matrix. (B) Macroscopic image of an explanted humanized ossicle. Scale bars represent 0.5 cm. (C) Hematoxylin and eosin (H&E) staining of a representative humanized ossicle. Left: overview of the ossicle structure; right: magnified view of the boxed region in the left panel. Scale bar, 100 µm. CB, cortical bone; TB, trabecular bone; HC, hematopoietic cells; a, adipocyte; v, vessel. (D) Immunohistochemical staining of the ossicle using an antibody against human mitochondria (brown). Red arrows indicate human mitochondria–positive cells. (E) Representative flow cytometry plots showing human CD45□ cells among viable cells isolated from ossicles eight weeks after intraossicle transplantation of 3 × 10□ human bone marrow CD34□ cells. (F) Quantification of the percentage of human CD45□ cells in ossicles (*n* = 8) compared with murine femurs 8 weeks after transplantation of CD34+ cells (*n* = 5). **: p < 0.01 (Mann-Whitney test). Data shown as individual data points (circles) and median (horizontal lines).

In the next step, human CD34□ hematopoietic bone marrow (BM) cells were injected intra-ossically (i.o) into 8-week ossicles. Human CD45□ cell engraftment within the ossicle was detectable as early as 4 weeks post-injection (Fig. S1A). By 8 weeks, stromal cells displaying an elongated, flattened morphology and positive human mitochondria staining were observed in the endosteal, peri-adipocytic, and interstitial regions. (Fig. S1B). Engraftment of human hematopoietic cells was significantly higher in the ossicles compared to the murine femurs (48.9% for ossicles vs. 2.3% for femurs), demonstrating the superior capacity of the humanized ossicle to support human hematopoiesis (Fig. 1E-F). Both human myeloid cells (CD15□, CD33□, CD66b□) and CD19□ lymphoid cells were detected, along with human CD34□ hematopoietic stem and progenitor cells (Fig. S1C-F), indicating that the humanized bone marrow microenvironment in NSG mice supports robust engraftment and differentiation of human hematopoietic cells.

### Transplantation of THPO overexpressing CD34+ bone marrow cells induces fibrosis in humanized ossicles

Thrombopoietin (THPO) overexpression has been shown to induce myelofibrosis in murine models [9, 18]. Humanized ossicles were therefore injected with human THPO-GFP bone marrow (BM) CD34□ cells and examined at 4, 5, 6, and 8 weeks after transplantation (i.o.). Staining for reticulin fibers was used as the gold standard to evaluate myelofibrosis using clinically-established criteria [19]. A relatively low grade of fibrosis was observed at 4 weeks after transplantation, whereas moderate to high-grade (grade 2-3) reticulin fiber deposition became evident at weeks 5 and 6 (Fig. S2A-B). By 8 weeks, a dense net of reticulin fibers was formed in the THPO ossicle marrow cavity, extending between hematopoietic areas and along trabecular bone surfaces, consistent with stable and well-established fibrosis (Fig. 2A). The median grade of week 8 ossicles engrafted with THPO-overexpressing CD34□ cells was grade 1.5 and was significantly higher compared to GFP controls (median grade 0) (Fig. 2B). Based on these findings, the 8-week time point was selected for subsequent studies and ossicles were assigned as 8-week huMF ossicles.

**Figure 2.**
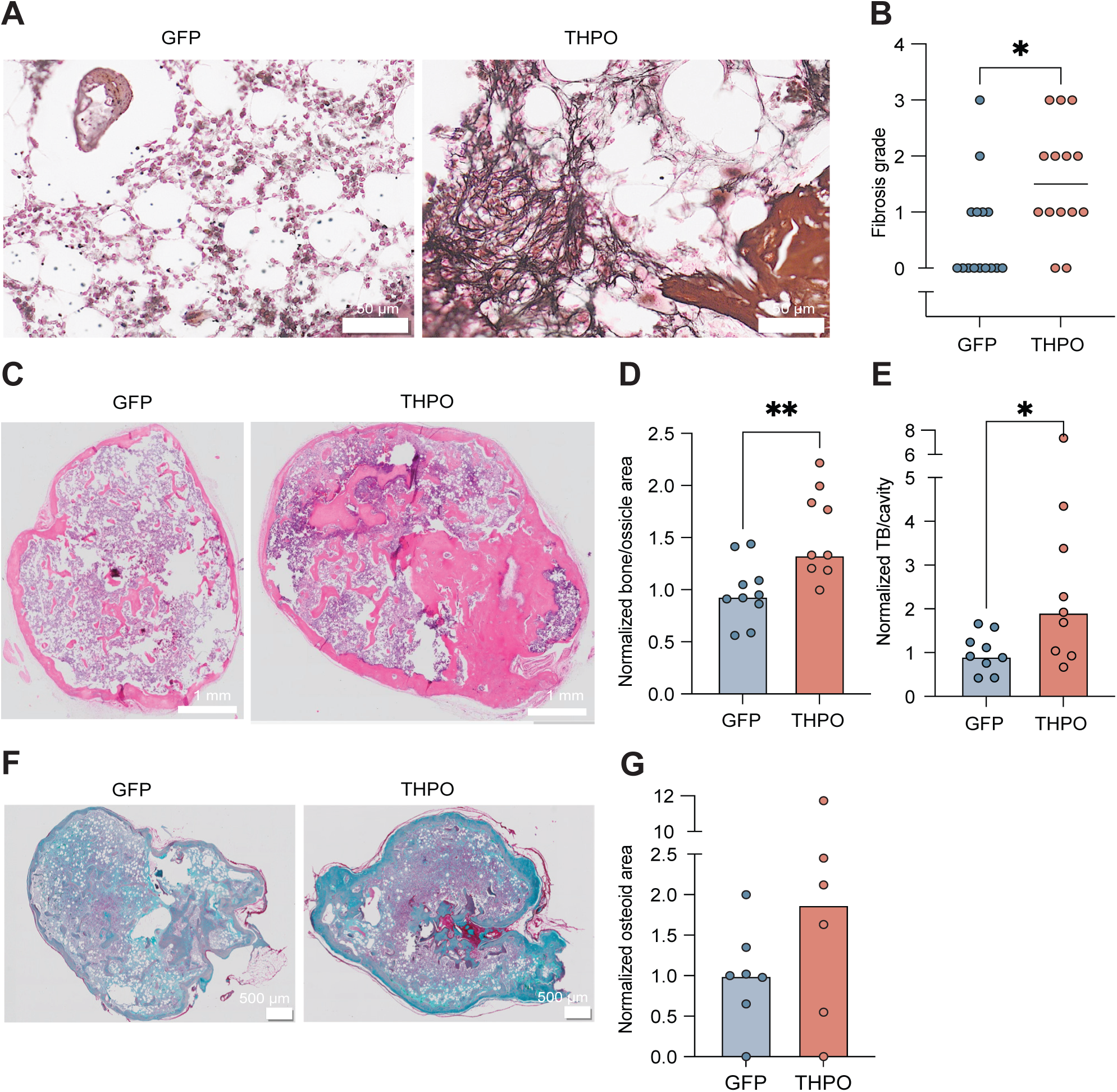
THPO overexpression induces fibrosis and osteosclerosis in huMF ossicles. (A) Representative images of Gordon and Sweet’s reticulin staining of GFP and THPO ossicles 8 weeks post-transplantation. Scale bar indicates 50 µm. (B) Blinded fibrosis grading was performed independently by two observers. Dot plot showing the fibrosis grade of individual ossicles (n = 15 for GFP and n = 14 for THPO). Statistical comparisons were performed using the Mann–Whitney test. *p< 0.05. (C) Representative H&E-stained images of GFP and THPO ossicles at 8 weeks post-transplantation. Scale bar: 1 mm. (D) Quantification of the normalized bone area to the total ossicle area in THPO ossicles relative to GFP controls (GFP: n = 10, THPO: n = 9; **p = 0.0001, Mann-Whitney test). (E) Quantification of the normalized trabecular bone (TB) to the ossicle cavity area relative to GFP controls (GFP: n = 8, THPO: n = 9; *p = 0.0244, Mann-Whitney test). (F) Representative RGB staining of GFP and THPO ossicles. Scale bar: 500 µm. (G) Quantification of the normalized osteoid area relative to GFP controls within the ossicle (GFP: n = 6, THPO: n = 6). Data shown as individual data points (circles) and median (horizontal lines).

### HuMF ossicles exhibit increased osteosclerosis

Another well-recognized feature of myelofibrosis is osteosclerosis. To investigate whether ossicles transplanted with THPO overexpressing CD34^+^ cells (HuMF ossicles) showed increased bone formation, high-resolution images from scanned H&E-stained slides were used to quantify the ossicle bone areas. Here, our data demonstrated that huMF ossicles had a higher bone-to-total ossicle area ratio compared to the GFP group, indicating increased bone formation caused by THPO-overexpressing CD34^+^ cells (Fig. 2C-D). Furthermore, huMF ossicles showed a significant increase in trabecular bone formation compared with controls (Fig. 2E).

We further performed trichrome staining (also known as RGB staining) to distinguish between different stages of osteogenesis based on the differences in collagen staining patterns between unmineralized bone and mineralized bone matrix. RGB staining demonstrated that cortical bones (outer layer) of the ossicles were mainly composed of the mineralized hardened matrix while the trabecular bones (the porous, lattice-like inner bone) were mostly composed of an unmineralized matrix. i.e. osteoid (Fig. 2F). Here, quantitative analysis demonstrated that huMF ossicles contained a higher percentage of unmineralized osteoid (purple) compared to GFP controls, suggesting a more pronounced remodelling (Fig. 2G).

Collectively, our results indicate that THPO-overexpressing hematopoietic cells promote pathological bone remodeling within the ossicle, recapitulating the osteosclerotic changes characteristic of myelofibrosis.

### Ossicles injected with THPO-overexpression CD34+ cells show a skewed myeloid/lymphoid ratio, increased megakaryopoiesis and pronounced megakaryocyte clustering

In vitro, THPO overexpression led to an increased percentage of CD41a□CD42b□ megakaryocytes, even in the absence of exogenous THPO in the culture medium (Fig. S3A-B). This result confirmed the functionality of lentiviral THPO overexpression in vitro. Flow cytometry analysis of ossicle cells showed that intraossicle (i.o) injection of both GFP-and THPO-overexpressing CD34□ cells resulted in a robust ossicle engraftment, with comparable levels of human CD45□ cells (Fig. 3A-B, S3C). Eight weeks following injection, the fraction of THPO-overexpressing cells was significantly increased compared with the initial frequency at the time of injection compared to controls (Fig. S3D), indicating a more rapid expansion of THPO-overexpressing cells and suggesting a proliferative and/or survival advantage conferred by THPO overexpression. Moreover, ossicles injected with THPO-overexpressing CD34□ cells exhibited myeloid skewing, as indicated by a higher ratio of myeloid (CD15□, CD33□, CD66b□) to CD19□ lymphoid cells (Fig. 3C-D).

**Figure 3.**
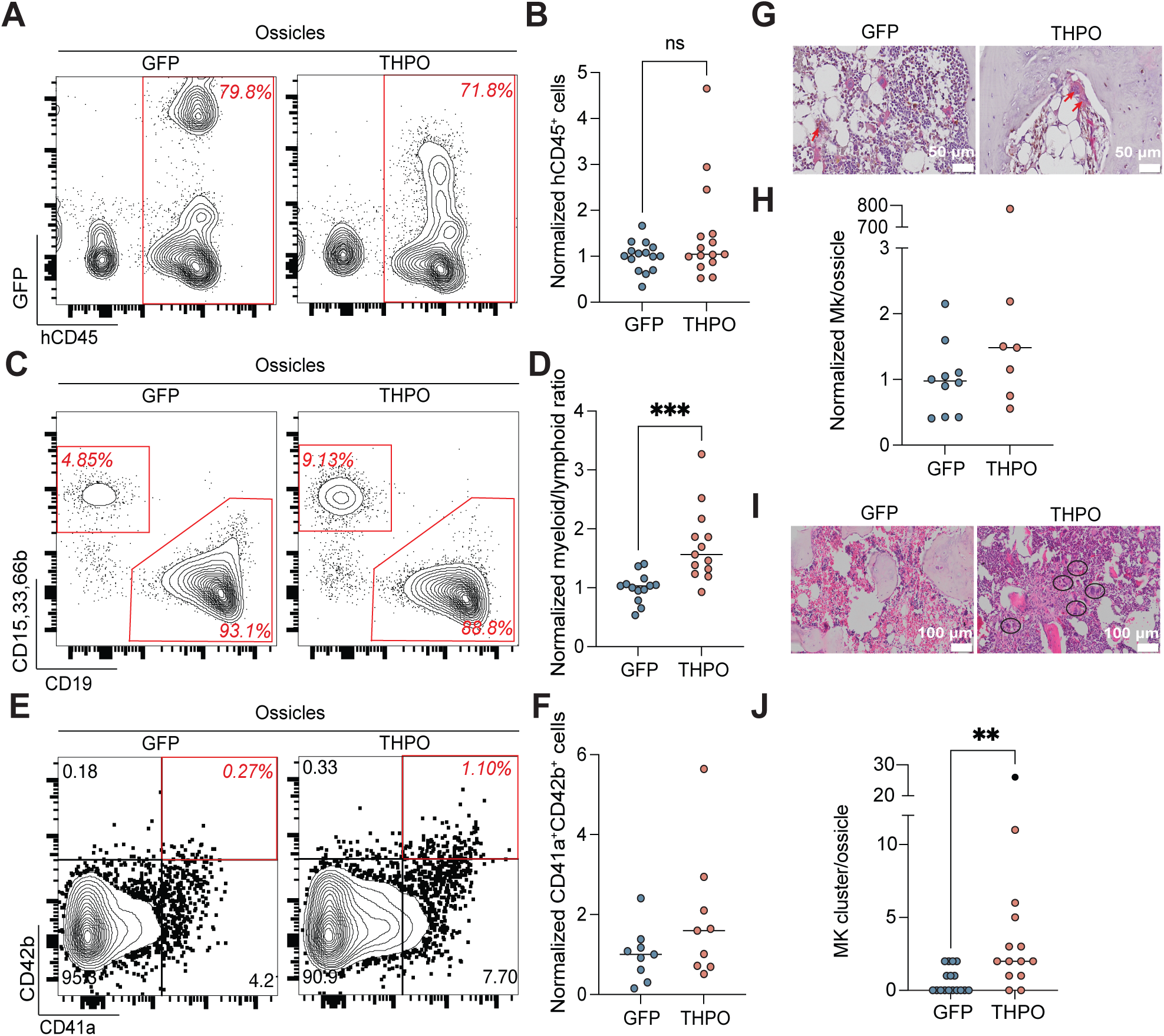
THPO overexpression induces a skewed myeloid/lymphoid ratio and megakaryocyte dysplasia in humanized ossicles. (A) Representative FACS plots showing human CD45□ cells in GFP and THPO ossicles 8 weeks after intraossicle transplantation. (B) Quantification of the normalized frequency of human CD45□ cells in THPO ossicles relative to GFP controls 8 weeks post-transplantation (GFP: n=15, THPO: n=15). (C) Representative FACS plots showing human CD15□CD33□CD66b□ cells versus CD19□ cells in GFP and THPO ossicles 8 weeks after intraossicle transplantation. (D) Quantification of the myeloid to lymphoid ratio in THPO ossicles relative to GFP controls 8 weeks post-transplantation (GFP: *n*=13, THPO: n=13). (E) Representative FACS plots (left and middle panels) showing human CD41a□ cells versus CD42b□ cells in GFP and THPO ossicles 8 weeks after intraossicle transplantation. (F) Quantification of the normalized frequency of human CD41a^+^CD42b^+^ cells in THPO ossicles relative to GFP controls 8 weeks post-transplantation (GFP: *n* =9, THPO: n=9). (G) Dual-color immunohistochemical (IHC) staining of ossicles showing human mitochondria (brown) and human CD61 (red). Red arrows indicate human megakaryocytes in GFP and THPO ossicles. Scale bar: 50 μm. (H) Quantification of megakaryocytes per ossicle. (I) H&E staining of GFP and THPO ossicles. Right panel: Megakaryocyte clusters are indicated by black circles. Scale bar: 100 μm. (J) Quantification of number of megakaryocyte clusters per ossicle. (**p = 0.0014, Mann-Whitney test). Data shown as individual data points (circles) and median (horizontal lines).

Additionally, 8-week huMF ossicles showed a tendency to increased frequencies of human CD41a^+^CD42b^+^ megakaryocytes compared to controls (Fig. 3E-F). However, differences were not statistically significant. In order to corroborate the flow cytometry findings, we quantified the number of human megakaryocytes in H&E-stained FFPE slides. IHC staining with human mitochondria marker showed an increased number of human megakaryocytes in huMF ossicles (Fig. 3G-H). As shown in Fig. 3H, there was a 1.5-fold increase in total megakaryocytes in huMF ossicles compared with GFP controls. We also found significantly increased megakaryocytic clustering in huMF ossicles (Fig. 3I-J).

### THPO overexpression triggers HSPC migration from ossicles to other hematopoietic organs in NSG mice

In 8-week huMF ossicle-bearing mice, we observed an increased percentage of human CD45□ hematopoietic cells in the femur and spleen compared to controls (Fig. 4A-D). Migrated human cells were primarily detected in the femurs, followed by the spleen and, to a much lesser extent, the liver (Fig. 4A-D, S4A-B). This distribution pattern likely reflects the superior hematopoietic-supportive microenvironment of the femur relative to the spleen and liver.

**Figure 4.**
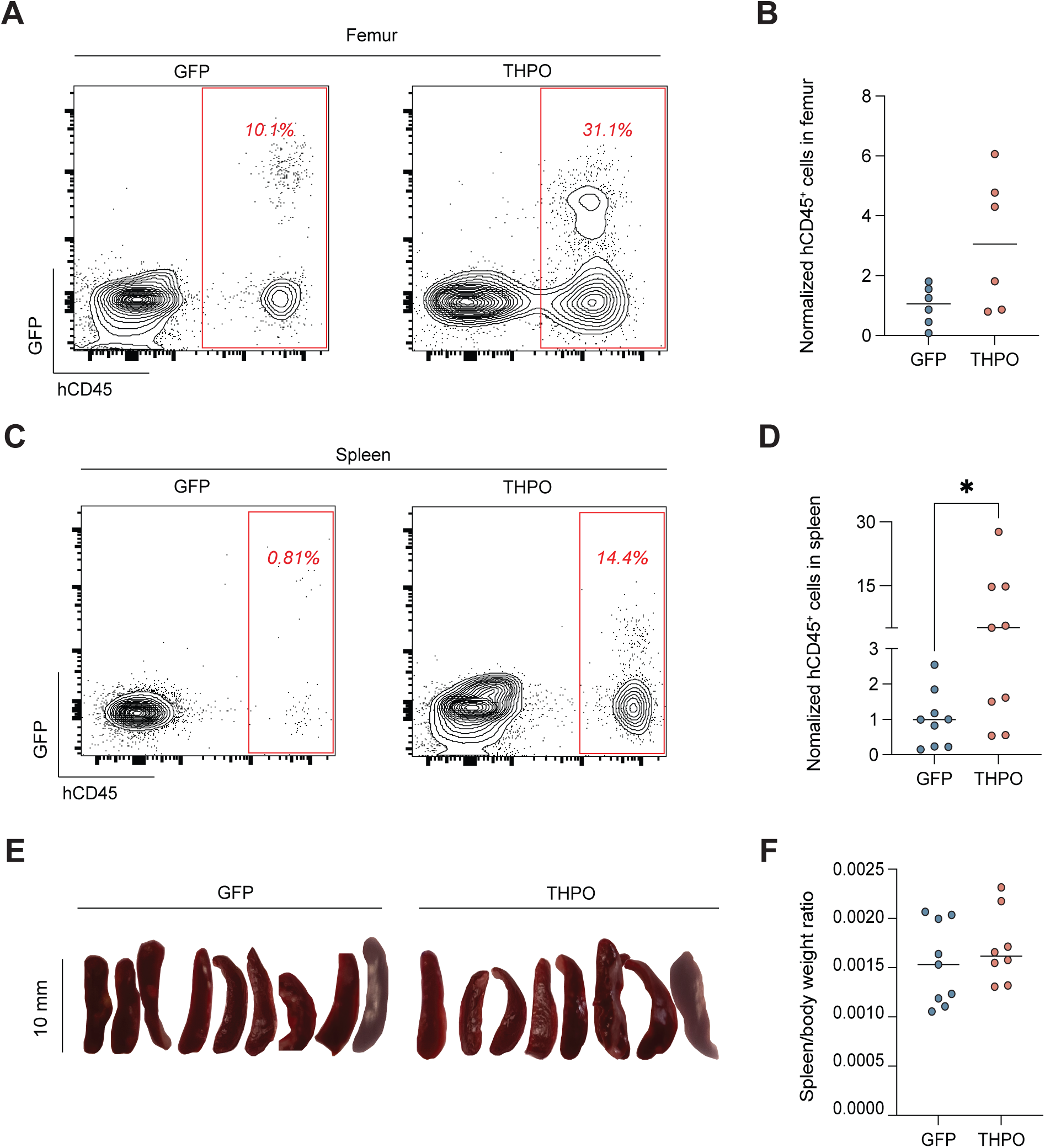
THPO overexpression induces migration of human hematopoietic cells from ossicles. (A) Representative FACS plots showing human CD45□ cells in femurs from GFP and THPO groups 8 weeks after intraossicle transplantation. (B) Quantification of the normalized frequency of human CD45□ cells in femurs of the THPO group relative to GFP controls at 8 weeks post-transplantation (n = 6 per group). (C) Representative FACS plots showing human CD45□ cells in spleens from GFP and THPO groups 8 weeks after intraossicle transplantation. (D) Quantification of the normalized frequency of human CD45□ cells in spleens of the THPO group relative to GFP controls at 8 weeks post-transplantation. (n = 9 per group. *p < 0.05 (Mann-Whitney test). (E) Macroscopic images of spleens from GFP and THPO ossicle-bearing mice. (F) Spleen-to-body weight ratio (GFP: n = 9, THPO: n = 8). Data shown as individual data points (circles) and median (horizontal lines).

A trend toward increased migration of human hematopoietic cells to the femur was observed in the THPO group (Fig. 4A-B), which however did not reach statistical significance. In contrast, significantly higher levels of human hematopoietic cells were detected in the spleens of the THPO group (Fig. 4C-D), despite comparable spleen sizes and spleen-to-body weight ratios of the THPO and GFP controls (Fig. 4E-F).

Together, these findings indicate an enhanced migration of human hematopoietic cells from humanized ossicles to suboptimal hematopoietic niches in the THPO group, suggesting that overexpression of THPO in CD34□ cells caused alterations of the ossicle hematopoietic microenvironment (HME) that favoured migration.

### SPP1/OPN is Upregulated in Myelofibrosis and SPP1 Antibody Treatment Partially Rescues MF Pathology

The enhanced bone formation observed in THPO ossicles suggested an increased activity of human osteoprogenitors within the humanized bone marrow microenvironment. Using single-cell RNA sequencing (scRNA-seq), we have in a previous study identified CD56□ cells as osteochondroprogenitors characterized by exclusive expression of secreted phosphoprotein I (SPP1)/osteopontin (OPN), which mediates their interactions with hematopoietic cells through an SPP1-dependent signaling axis [20]. Based on these findings, we performed immunostaining for CD56 and SPP1/OPN to assess osteoprogenitor activation in THPO ossicles. Immunofluorescence staining indicated a higher level of SPP1/OPN expression in THPO ossicles compared with GFP controls (Fig. 5A), suggesting enhanced osteoprogenitor activity and matrix production. These findings were confirmed in clinical bone marrow biopsies from patients with primary MF compared with hematologically healthy controls (Fig. 5B). Consistent with our huMF ossicle findings, patient MF samples exhibited significantly higher expression of SPP1/OPN (median SPP1 intensity: 3.38, n = 3) compared with controls (median SPP1 intensity: 2.71, n = 3).

**Figure 5.**
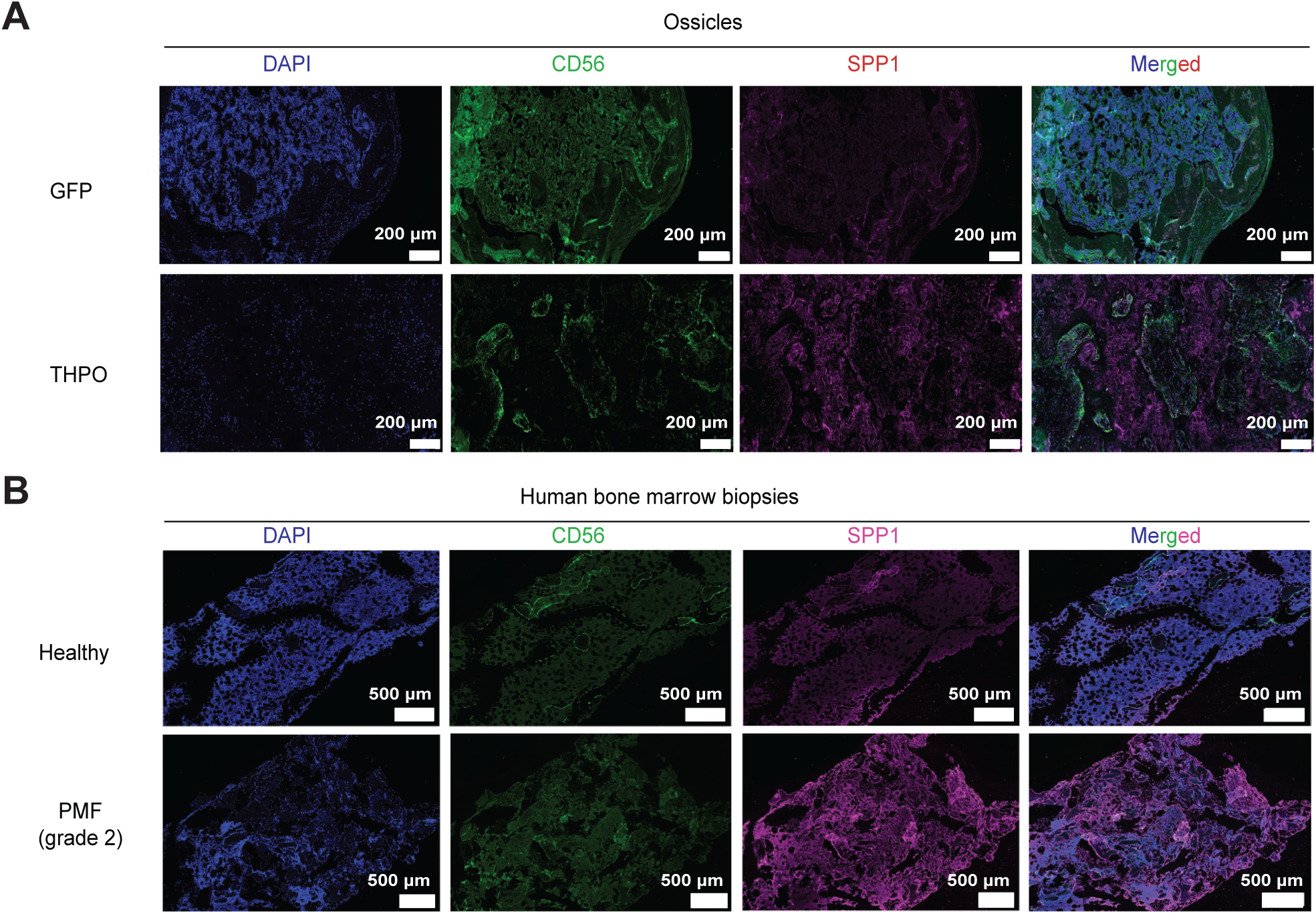
Increased SPP1 expression in huMF ossicles and PMF patient bone marrows. Representative images of CD56 and SPP1 immunostaining in GFP and THPO ossicles (A) and in bone marrow biopsies from healthy donors and patients with primary myelofibrosis (PMF) (B). Single channel images showing Nuclei stained with DAPI (blue), and antibody staining with CD56-AF488 (green) and SPP1-AF647 (red).

In summary, these findings demonstrate that THPO overexpression in CD34□ cells promotes enhanced osteogenic remodeling within the ossicle, potentially driven by the activation of CD56□ osteoprogenitors, which may contribute to niche remodeling through their capacity to produce SPP1/OPN. Consistent with this, SPP1/OPN expression was also found to be elevated in bone marrow biopsies from PMF patients, supporting the clinical relevance of this pathway.

To test the functional relevance of SPP1/OPN overexpression, we then went on to treat huMF ossicle mice with anti-SPP1 antibody (Fig. 6A). Histological analysis revealed a marked reduction in fibrosis in the hMF ossicles following anti-SPP1 treatment (Fig. 6B-C) and there was a trend toward reduced bone area (Fig. 6D-E). Even though the engraftment of human hematopoietic cells in ossicles (Fig. 6F), spleen (Fig. 6G) and femurs (Fig. 6H) was not affected, anti-SPP1 treatment attenuated the increased myeloid-to-lymphoid ratio and reduced CD41□ megakaryocyte frequencies (Fig. 6I).

**Figure 6.**
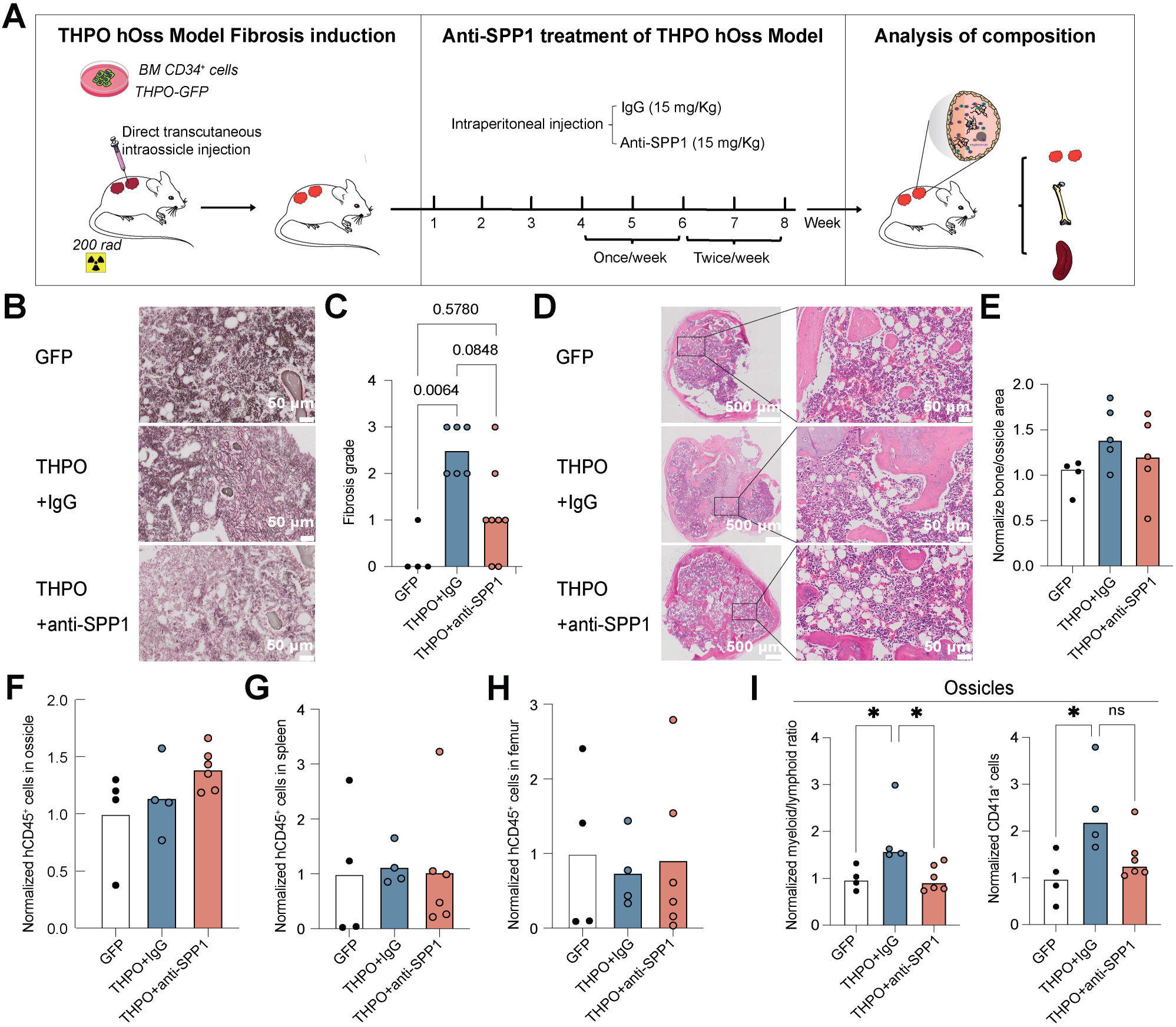
Anti-SPP1 neutralizing antibody treatment reduces fibrosis and attenuates the skewed myeloid-to-lymphoid ratio in huMF ossicles. (A) Schematic overview of the experimental design. Four weeks after intra-ossicle transplantation of THPO-overexpressing CD34□ cells, mice were treated with either anti-SPP1 or an isotype control antibody (both at 15 mg/kg) once weekly for two weeks. Treatment was subsequently increased to twice weekly for an additional two weeks. Mice were sacrificed 8 weeks after intra-ossicle injection, and ossicles were analyzed for fibrosis, osteosclerosis, and human hematopoietic cell engraftment. hOss, humanized ossicle. (B) Representative images of Gordon and Sweet’s reticulin staining of GFP and THPO ossicles from mice treated with either anti-SPP1 antibody or IgG control 8 weeks post-transplantation. Scale bar indicates 50 µm. (C) Quantification of fibrosis grades in THPO ossicles after treatment with anti-SPP1 antibody or IgG control, compared with GFP controls. P values are indicated on top of comparison bars. (D) Hematoxylin and eosin (H&E) staining of the representative GFP and THPO ossicles from mice treated with either anti-SPP1 antibody or IgG control group. Left panel: Overview of the ossicle, with selected regions of interest indicated by frames. Scale bar, 500 µm. Right panel: Blow-up images corresponding to the framed areas in the overview, highlighting detailed tissue architecture. Scale bar, 50 µm. (E) Quantification of the normalized bone-to-ossicle area ratio in THPO ossicles following treatment with anti-SPP1 antibody or IgG control, relative to GFP controls. (F) Quantification of the normalized human CD45+ cells in THPO ossicles after treatment with anti-SPP1 antibody or IgG control, compared with GFP controls. Quantification of the percentage of human CD45□ cells in the spleens (G) and femur (H) of THPO ossicle–bearing mice following treatment with anti-SPP1 antibody or IgG control, compared with GFP ossicle–bearing mice. (I) Quantification of the normalized myeloid-to-lymphoid ratio in THPO ossicles following treatment with anti-SPP1 antibody or IgG control, relative to GFP controls (left). Quantification of the normalized percentage of CD41□ cells in THPO ossicles following treatment with anti-SPP1 antibody or IgG control, relative to GFP controls (right). *p < 0.05 (Kruskal-Wallis test).

## Discussion

A number of experimental models have been developed to study myelofibrosis pathogenesis. Murine models based on genetic modification of hematopoiesis including transgenic or retroviral overexpression of THPO [18], JAK2^V617F^ [22], MPL^W515L^ [7], and CALR mutations [23] recapitulate key disease features. However, because fibrosis develops within an entirely murine microenvironment, these models do not fully capture species-specific stromal responses and niche–hematopoietic cell interactions relevant to the human disease. To address species differences in hematopoiesis, xenotransplantation models using human mutated HSPCs have been widely adopted. In these models, human CD34□ cells carrying driver mutations are transplanted into immunodeficient mice to study human clonal hematopoiesis in vivo [17, 24]. Although these systems capture cell-intrinsic effects of disease mutations, fibrosis development is often limited, likely due to lacking interactions between human hematopoietic cells and murine stromal cells and potential differences in fibrogenic pathways.

As an alternative, recent advances in stem cell biology have enabled the development of in vitro fibrosis models based on induced pluripotent stem cell (iPSC)-derived organoids [25]. These systems allow a controlled reconstruction of hematopoietic and stromal components, and they are particularly valuable for mechanistic studies and high-throughput drug screening. However, the in vitro nature of this approach limits their ability to fully capture the spatial organization, vascularization, mechanical forces, and, importantly, the long-term remodeling characteristics of bone marrow fibrosis development.

In this context, humanized bone marrow niche models, including ectopic ossicle systems, offer a unique intermediate approach for modeling myelofibrosis in vivo. Humanized bone marrow tissues established in immunodeficient mice enable sustained engraftment of human hematopoietic cells and, thus, allow to study pathological BM remodeling within a species-matched niche.

Accordingly, we herein demonstrate an ossicle-based system that recapitulated key features of myelofibrosis and allowed for a direct investigation of human hematopoietic-stromal crosstalk and therapeutic targeting of niche-derived factors.

We established a humanized bone marrow microenvironment (hu ossicles) in immunodeficient NSG mice by transplantation of human BM stromal cells and showed that overexpression of THPO in human CD34□ cells was sufficient to induce fibrotic ossicles (huMF ossicles). These huMF ossicles showed key hallmarks of myelofibrosis, including myeloid skewing, megakaryocytic dysplasia, progressive reticulin fibrosis, extramedullary hematopoiesis, and pathological bone remodeling.

Importantly, we identify SPP1/OPN as a critical mediator of the pathological changes and showed that SPP1/OPN neutralization partially rescued the myelofibrotic phenotype. Clearly, this finding indicates that the anti-SPP1 antibody approach might present a possible novel therapy for myelofibrosis, which needs to be addressed in future studies.

In the murine system, THPO overexpression leads to rapid and consistent induction of bone marrow fibrosis. We therefore chose to employ THPO overexpression in human CD34+ hematopoietic cells as a first step to induce fibroses in humanized ossicles. Certainly, alternative approaches including generation of huMF ossicles using mutated hematopoietic cells might be possible and will be evaluated in future experiments.

Consistent with murine models of myelofibrosis [18, 26, 27], we saw a trend toward expansion and clustering of megakaryocytes in the THPO ossicles, which is a defining feature of myelofibrosis pathogenesis. Progressive ossicle reticulin fibrosis developed over time and was well-established eight weeks post-transplantation of THPO-overexpressing CD34□ cells. Notably, the spatial distribution and architecture of reticulin fibers in the MF ossicles closely resembled those observed in human myelofibrosis, with dense networks extending between hematopoietic areas and along trabecular bone surfaces. These findings demonstrate that dysregulated THPO signaling in human hematopoietic cells is sufficient to drive fibrotic remodeling within a human stromal context, supporting the notion that hematopoietic cell–intrinsic alterations can initiate and sustain fibrotic niche remodeling.

In myelofibrosis patients, the bone marrow’s ability to support hematopoiesis is compromised, leading to a shift of blood cell production to other hematopoietic organs. Consequently, a hallmark feature of myelofibrosis is extramedullary hematopoiesis, often manifested as splenomegaly and hepatomegaly. In parallel with fibrotic progression, we observed enhanced migration of human hematopoietic cells from the humanized ossicles to murine hematopoietic organs, most prominently the spleen. This redistribution mirrors extramedullary hematopoiesis. The relatively low proportion of human hematopoietic cells detected in the spleen may account for the absence of overt splenomegaly at this stage and splenomegaly might develop at later stages as hematopoietic cell accumulation progressively increases, which we however have not investigated yet.

Quantitative analysis revealed increased human CD45□ cell engraftment in both the murine femur and spleen of THPO ossicle–bearing mice compared with controls. This difference likely reflects intrinsic variations in the hematopoietic-supportive capacity of these organs. While the murine femur represents a relatively permissive niche for human hematopoietic cells even under baseline conditions, the spleen acts as a more responsive extramedullary site where pathological cues may drive a more pronounced accumulation of human cells.

Importantly, the preferential accumulation of human hematopoietic cells in the spleen is consistent with a key clinical hallmark of myelofibrosis, which is the mobilization of HSPCs from the BM and their expansion in extramedullary organs. The absence of statistical significance in femoral engraftment does not exclude biologically meaningful migration but rather suggests that niche occupancy or saturation may limit further accumulation.

Osteosclerosis represents another hallmark of myelofibrosis and remains incompletely understood mechanistically. Our data demonstrate that THPO overexpression induced excessive bone formation within the ossicles, characterized by increased trabecular bone area and accumulation of unmineralized osteoid. These findings suggest an imbalance between bone formation and maturation, indicative of active but dysregulated osteogenesis. The presence of abundant osteoid further implies ongoing microenvironmental remodeling rather than static bone deposition.

Mechanistically, we identified a marked upregulation of CD56 and SPP1/OPN in THPO ossicles, implicating activation of osteochondroprogenitor populations [20]. In line with our previous single-cell transcriptomic analyses, CD56□ cells expressing high levels of SPP1 appear to represent a specialized osteoprogenitor subset that engages in bidirectional crosstalk with hematopoietic cells [20]. The concordant upregulation of CD56 and SPP1 in bone marrow biopsies from primary myelofibrosis patients underscores the clinical relevance of these findings and supports the translational validity of our model. These observations are consistent with recent studies demonstrating that fibrosis-inducing hematopoietic cells can reprogram bone marrow stromal progenitors toward a profibrotic and osteogenic state. In particular, peritrabecular osteolineage cells marked by NCAM1/CD56 have been shown to expand during myelofibrosis and contribute to osteosclerosis through activation of Wnt/β-catenin–dependent programs [28]. The expansion of CD56□ stromal populations observed in our THPO-driven model therefore likely reflects a similar injury-induced stromal remodeling process, in which osteolineage progenitors are activated and progressively occupy the central marrow space. Moreover, SPP1/OPN has emerged as a central profibrotic mediator in myelofibrosis, promoting stromal cell activation and collagen production, correlating with fibrosis severity, and representing a promising therapeutic target [14, 29].

Importantly, therapeutic neutralization of SPP1/OPN attenuated several THPO-induced pathological features, including myeloid skewing, megakaryocyte expansion, and fibrosis severity. Although reductions in splenic engraftment and bone area did not reach statistical significance, the observed trends suggest that longer treatment duration or combination approaches may further enhance therapeutic efficacy, which can be studied in future experiments. Our findings thus strongly suggest SPP1/OPN not merely as a biomarker of disease-associated stromal activation, but as a functional mediator of hematopoietic–stromal dysregulation in myelofibrosis.

A key limitation of this model is that it does not fully recapitulate the native bone marrow environment. Ossicles are ectopically formed and lack full anatomical integration with native bone marrow, resulting in incomplete exposure to physiological cues such as mechanical loading. Moreover, key microenvironmental components, including vasculature and neural elements, remain of murine origin, potentially influencing niche behavior and limiting the faithful recapitulation of human bone marrow physiology.

Despite these limitations, our study establishes a humanized in vivo platform that faithfully models key aspects of human myelofibrosis and reveals a central role for SPP1/OPN-driven osteogenic and fibrotic remodeling downstream of aberrant THPO signaling. Targeting the SPP1 axis may therefore represent a promising therapeutic strategy aimed at restoring bone marrow niche function in myelofibrosis, complementing current treatments that primarily target malignant hematopoietic clones.

## Notes

### Competing Interest Statement

The authors have declared no competing interest.

### Summary of Updates

The manuscript has been substantially revised to improve clarity and incorporate new data. The main figures have been reorganized, and additional data have been included in both the main and supplementary figures. A new supplementary table (Table S1) has also been added. The main text has been updated accordingly to reflect these changes.

### Funding

- Swedish Childhood Cancer Foundation
- Swedish Cancer Foundation
- Swedish Bloodcancer Association (Blodcancerförbundet)
- Foundation Siv-Inger and Per-Erik Anderssons minnesfond
- John Persson Foundation
- Avtal om Läkarutbildning och Forskning (Government Public Health Grant)

