## Supplementary material for "A humanized ossicle model of myelofibrosis reveals THPO-driven fibrosis, osteosclerosis and SPP1-dependent microenvironmental remodeling": FigS1

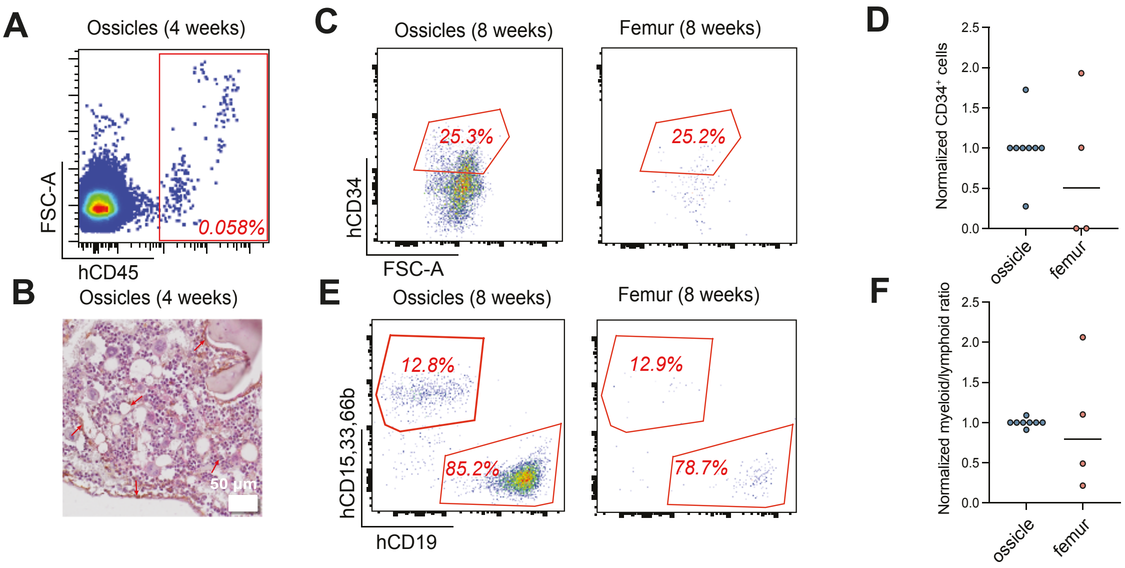


**Figure S1. Human hematopoietic engraftment and lineage differentiation within the humanized ossicle.** (A) Immunohistochemical staining of ossicle sections using antibodies against human mitochondria (brown) and CD61 (red), confirming the presence of human stromal and megakaryocytes, respectively. Red arrows indicate human mitochondria–positive stromal cells. (B) Representative flow cytometry (FACS) plots showing human $CD45^{+}$ cells in the humanized ossicle 4 weeks after intraossicle transplantation of human $CD34^{+}$ cells. (C) Representative FACS plots showing human $CD34^{+}$ hematopoietic stem and progenitor cells in the ossicle 8 weeks after transplantation in ossicle and femur. (D) Quantification of the fold change of the percentage of $CD34^{+}$ cells in ossicles and femurs 8 weeks after intraossicle transplantation of bone marrow $CD34^{+}$ cells. (E) Representative FACS plots showing lineage distribution of human hematopoietic cells in the ossicle 8 weeks after transplantation, including human $CD19^{+}$ lymphoid cells and $CD15^{+}CD33^{+}CD66b^{+}$ myeloid cells. (F) Quantification of the fold change of myeloid-to-lymphoid ratio in ossicles and femurs 8 weeks after intraossicle transplantation of bone marrow $CD34^{+}$ cells.
