## Supplementary material for "A humanized ossicle model of myelofibrosis reveals THPO-driven fibrosis, osteosclerosis and SPP1-dependent microenvironmental remodeling": FigS2

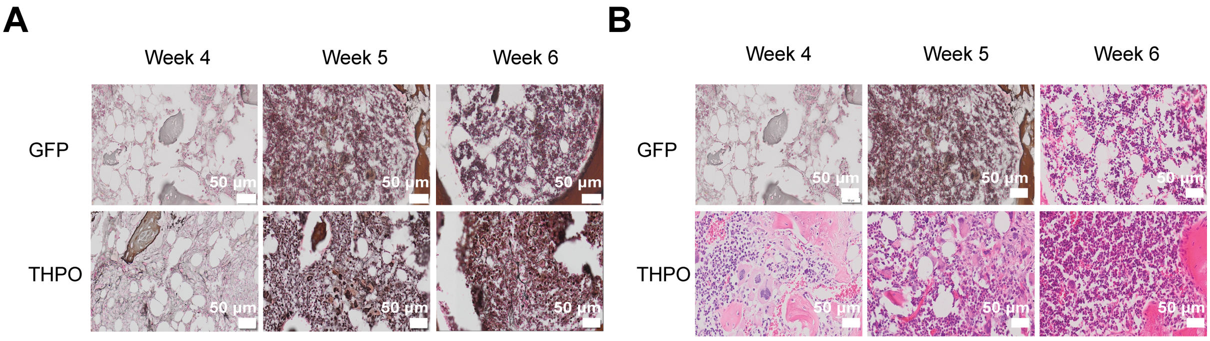


**Figure S2.** Representative Gordon and Sweet's reticulin staining (A) and H&E staining (B) of GFP and THPO ossicles at 4, 5, and 6 weeks post-transplantation.
