## Supplementary material for "A humanized ossicle model of myelofibrosis reveals THPO-driven fibrosis, osteosclerosis and SPP1-dependent microenvironmental remodeling": FigS3

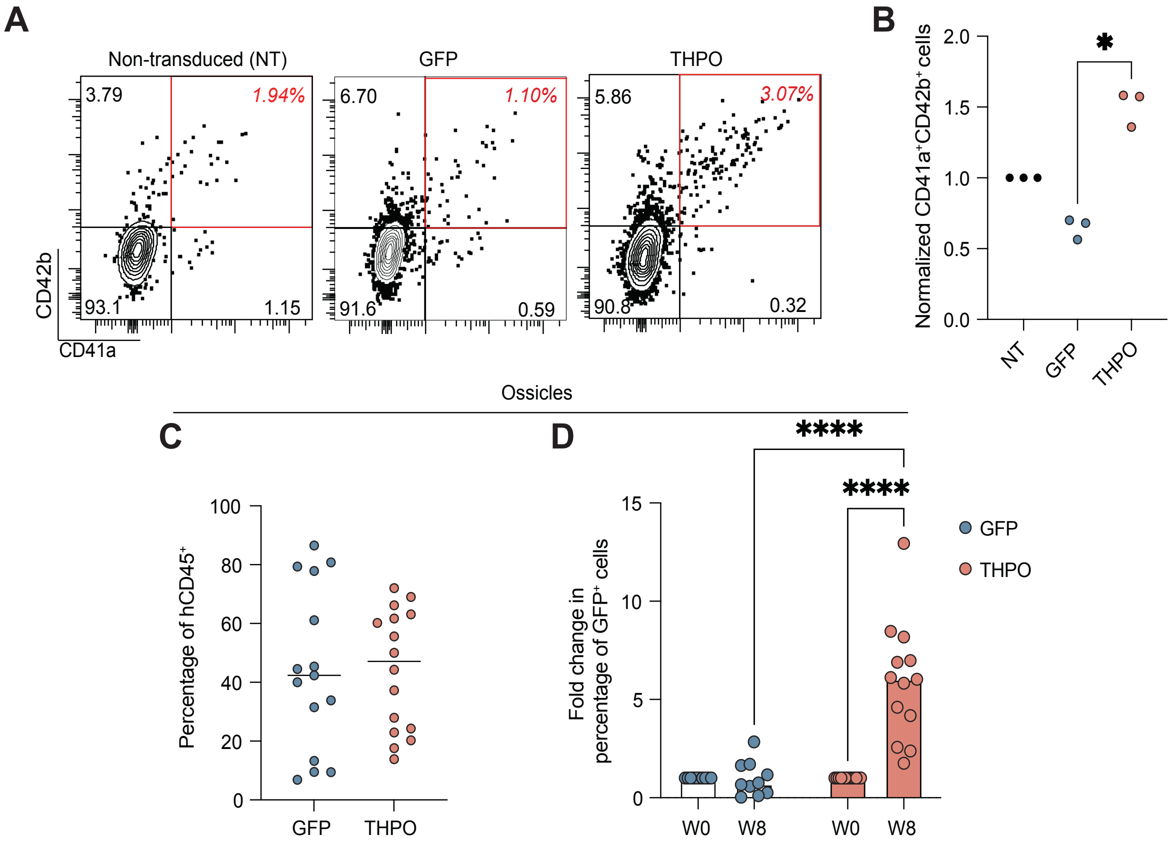


**Figure S3.** (A) Representative FACS plots showing CD41a$^+$CD42b$^+$ cells generated from BM-derived CD34$^+$ cells cultured for 5 days in SFEM medium supplemented with SCF (25 ng/mL), IL-3 (10 ng/mL), and IL-6 (10 ng/mL). (B) Quantification of the fold change in the percentage of CD41a$^+$CD42b$^+$ cells relative to non-transduced (NT) controls after 5 days of culture (mean ± SD, n = 3; $p < 0.05$, Mann–Whitney test). (C) Quantification of the percentage of human CD45$^+$ cells in THPO versus GFP ossicles 8 weeks after transplantation of CD34$^+$ cells. (D) Quantification of the fold change in the percentage of GFP$^+$ cells before transplantation (week 0, w0) and 8 weeks after transplantation (week 8, w8). Data are shown as indivudal data points (circles) and median (bars). (GFP: n = 10, THPO: n = 13, ****$p < 0.0001$, Two-way ANOVA test).
