## Supplementary material for "A humanized ossicle model of myelofibrosis reveals THPO-driven fibrosis, osteosclerosis and SPP1-dependent microenvironmental remodeling": FigS4

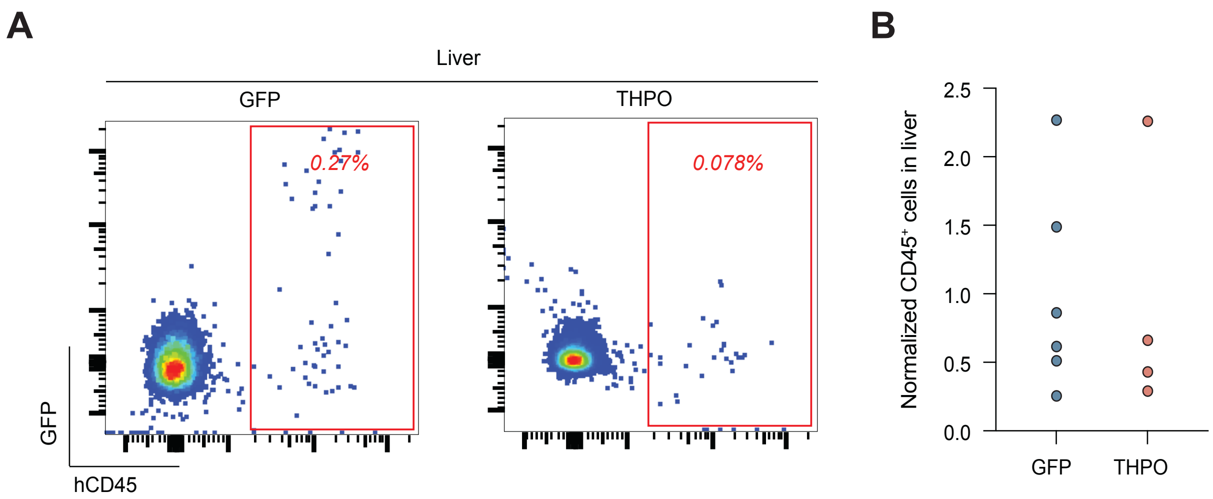


**Figure S4.** (A) Representative flow cytometry (FACS) plots showing human CD45$^{+}$ cells in the liver of ossicle-bearing mice 8 weeks after intra-ossicle transplantation of human CD34$^{+}$ cells. (B) Quantification of the normalized frequency of human CD45$^{+}$ cells in the liver of ossicle-bearing mice relative to GFP controls.
