## Supplementary material for "A humanized ossicle model of myelofibrosis reveals THPO-driven fibrosis, osteosclerosis and SPP1-dependent microenvironmental remodeling": Table S1

Table S1. Patient characteristics.

| PMF patients and normal controls | | | | | |
| --- | --- | --- | --- | --- | --- |
| Sample ID | Diagnosis | | Fibrosis grade | Sex | Age |
| P1 | PMF |  | 2 | F | 69 |
| P2 | PMF |  | 2 | F | 32 |
| P3 | PMF |  | 2 | M | 46 |
| C1 | Normal control | | 0 | M | 76 |
| C2 | Normal control | | 0 | M | 43 |
| C3 | Normal control | | 0 | M | 49 |

Abbreviations: PMF, primary myelofibrosis; P, patient; C, control; M, male; F, female.
